# Electroconvulsive stimuli reverse neuro-inflammation and behavioral deficits in a mouse model of depression

**DOI:** 10.1101/2023.05.13.540577

**Authors:** Alasdair G Rooney, Alastair M Kilpatrick, Charles ffrench-Constant

**Affiliations:** Centre for Regenerative Medicine Institute for Regeneration and Repair The University of Edinburgh Edinburgh Bioquarter 5 Little France Drive Edinburgh EH16 4UU; Norwich Medical School University of East Anglia Norwich Research Park Norwich NR4 7TJ

**Keywords:** Electroconvulsive, depression, mouse, SGZ, microglia, neuro-inflammation

## Abstract

**Background:** Electroconvulsive therapy is a fast, safe, and effective treatment for severe clinical depression but there is an ongoing search for mechanistic insights.

**Methods:** We used a mouse neuro-endocrine model of depression to examine behavioral, cellular, and molecular effects of electroconvulsive stimuli (ECS).

**Results:** The behavioral response to repeated ECS correlated with adult neurogenesis, more strongly in the ventral than dorsal hippocampus. Subsequent RNA-seq analysis targeting the ventral subgranular zone (SGZ) delineated ECS-responsive molecular pathways that were shared between naive and depressive-state conditions, and which may represent core biological responses to seizure induction. Other pathways responded to ECS preferentially in the depressive state, suggesting further state-specific mechanisms. By comparing gene set pathways reciprocally altered in depressed-state animals then reversed by ECS, we identified and validated neuro-inflammation as a candidate regulator of the antidepressant response. We further identified 56 novel candidate ‘antidepressant response’ genes in the ventral SGZ that may contribute to recovery, half of which have been implicated in human neuropsychiatric phenotypes.

**Conclusions:** Electroconvulsive stimuli reverse neuro-inflammation in a mouse model of depression. The results offer a detailed molecular characterization of potential SGZ antidepressant response-specific genes and pathways in brain regions implicated in depression.

## INTRODUCTION

Electroconvulsive therapy (ECT) is a highly effective treatment for severe depression, (1,2) but mechanisms of its antidepressant effect remain unclear. (3) Many preclinical studies have therefore examined the animal model of electroconvulsive shock (ECS) for cellular and molecular insights. (4) However, most of these studies delivered ECS to naive unstressed animals rather than first modeling depression. (5) This is a potentially important omission. ECT is only given to patients with significant mental disorders such as depression, so first modeling depression in animals **-** for example by manipulating the hypothalamic-pituitary axis (6) - may improve experimental validity. (7) It is also possible that the molecular and cellular mechanisms triggered in response to ECS in model states differ from those induced in naive or unstressed states. Modeling the disordered state may uncover translationally relevant mechanisms of action that are not present in naive conditions.

Here we replicate a preclinical model of depression by the chronic administration of Corticosterone (CORT) and confirm that ECS reverses CORT-induced depressive-like behaviors. Observing that the behavioral response to ECS correlates most strongly with ventral hippocampal neurogenesis, we perform RNA-sequencing analysis of the response of the ventral hippocampal niche to ECS, with and without prior CORT treatment. This analysis uncovers a reciprocal effect of CORT and ECS on neuro-inflammation and reveals novel ‘antidepressant response’ genes and gene pathways that may contribute to recovery during antidepressant treatment.

## METHODS AND MATERIALS

### Animals

Wild-type C57Bl/6 male mice were used beginning at age 8 weeks. Animals were housed in groups of 3 or more in individually ventilated cages under a 12hr light/dark cycle with ad libitum access to food and water.

### Experimental design

Two experimental cohorts of animals are reported in this paper. The experimental design for behavioral and cellular level analyses is illustrated in **Figure S1.** The design for molecular (laser-capture micro-dissection and RNA sequencing) analyses is illustrated in **Figure S2.**

### Corticosterone and EdU administration

Experimental animals received Corticosterone (CORT, 35 micrograms/ml, Cayman Chemical 16063) in autoclaved drinking water for a total of seven weeks. To fate-label proliferating cells, EdU (5-Ethynyl-2’-Deoxyuridine, Carbosynth NE08701, 10mg/ml) was administered on each day of ECS, by IP injection (0.1ml/mouse) while the animal was under the anesthetic for ECS.

### Electroconvulsive stimuli

Animals were anesthetized with isoflurane and O2 running at 4L/min for 60 seconds. They were rapidly positioned in ear clips attached to a custom-built ECS unit (kind gift from Dr. C.A. Stewart, University of Dundee). The electrical stimulus was delivered (1 sec duration, 150V, 25mA). Mice were closely observed for seizure activity. All mice receiving ECS had motor seizures lasting >10 and <30 seconds. Mice were moved to a recovery chamber until awake and fully ambulant (<5 minutes). Control animals were handled and treated in exactly the same way including anesthesia and positioning in ear clips, but minus delivery of the shock.

Morbidity and mortality from the procedure was zero.

### Behavioral tests

See Supplemental Methods.

### Tissue preparation

Animals were deeply anesthetized with an IP injection of medetomidate (1mg/kg, Orion Pharma) and ketamine (75mg/kg, Zoetis). They were then transcardially perfused with chilled 0.9% NaCl followed by 10ml 0.37% PFA (Sigma P6148) made up in 0.9% NaCl.

Brains were excised and stored in 4% PFA for 24h at 4C, cryoprotected in 15% and then 30% sucrose solution until sunk, and embedded in OCT (Cell Path, KMA-0100-00A). Brain blocks were stored at -80C until further use.

### Immunofluorescence

Floating sections were blocked for one hour with 10% NDS (Millipore S30) in 0.3% Triton X-100 (Fisher Scientific BP151-500) and 1xPBS (Gibco 7001-036, diluted to 1X in distilled water). Primary antibodies were incubated in blocking solution for 48h at 4C. Sections were washed x3 in PBS for a total of three hours. Secondary antibodies were incubated in blocking buffer for 24h at 4C. Sections were washed again, counterstained with Hoechst 33342 (Thermo Fisher Scientific 62249), and mounted to slides with Fluoromount-G (SouthernBiotech 0100-01).

### Correlations between dorsoventral neurogenesis and behavior

For each animal we calculated the ratio of their post-ECS to their pre-ECS performance. This gave a corrected value for each animal’s behavioral response to ECS relative to their own baseline. We plotted that value against the animal’s neurogenesis response in dorsal and ventral hippocampus. Linear regression was used to test whether the regression slope was significantly different from zero.

### Laser capture microscopy

Mice following the experimental design illustrated in **Figure S2** were killed at the appropriate time-points by cervical dislocation. Brains were quickly dissected and coronally bisected into anterior and posterior blocks. These blocks were immediately snap-frozen on liquid nitrogen and stored at -80°C until sectioning. For sectioning and laser-capture methodology see Supplemental Methods.

### RNA extraction and sequencing

In a preliminary test of four different RNA extraction methods (FFPE [Quiagen 73504], Microkit [Quiagen 74004], miRNeasy FFPE [Quiagen 217504], and Masterpure [MCR85102]) we obtained the best balance of quality and yield using the Quiagen FFPE kit [Quiagen 217504] (data not shown). RNA was extracted from micro-dissected tissue using this kit according to manufacturer instructions, omitting deparaffinisation but including Proteinase K and DNase steps. RNA was sequenced by SourceBioScience (Nottingham, UK) using an S1 flow-cell on an Illumina NovaSeq6000 sequencer with a target 60M, 50bp paired-end read depth.

### RNA-seq analysis

Transcript level counts were collapsed to gene level counts using a transcript to gene mapping table generated from Ensembl transcript and gene Ids; all analysis was carried out at gene level. Ensembl gene Ids were mapped to MGI gene symbols using the org.Mm.eg.db Bioconductor package (v.3.10.0). Differential expression was computed using the DESeq2 Bioconductor package (v.1.26.0). (8) For heat-map visualization of RNA-seq data, read counts were normalized with respect to library size using the regularized log (rlog) transform. The RNA-seq data associated with this study has been deposited at NCBI GEO (accession number GSE155706).

### Internal validation and interrogation of RNA-seq analysis

Internal validation was conducted using principal component analysis (PCA), heat-map visualization with hierarchical clustering, and intersection of differential expression results with known SGZ cell-type specific genes. To construct this SGZ cell-type specific gene list for the latter two approaches we combined gene lists of individual cell types previously identified in three published single-cell RNA-seq analyses of the SGZ niche (**Table S1**). (9–11)

### Pathway analysis pipeline

See Supplemental Methods.

### Independent validation of RNA-seq analysis

An independent cohort of animals was treated with CORT or VEH for 5 weeks, then each stream received ECS or Sham-ECS. Animals in this experiment were cage-mates of animals in the source RNA-seq experiment: their tissue was used for validation and not RNA extraction. These animals were sacrificed by perfusion fixation 48 hours after the final ECS, and brains processed for immunofluorescence as above.

### Ethical Statement

All experiments were pre-approved by University of Edinburgh veterinary staff and conducted in accordance with institutional and national guidelines.

## RESULTS

### Repeated ECS reverses depressive-like behavior

To establish a preclinical model of depression we utilized the chronic administration of corticosterone (CORT), which induces rodents to display reduced self-care, and increased anxiety. (12–15) As previously observed (16) CORT induced a depressive-like state evidenced by a poorer coat state (median coat state score VEH=4 [IQR 3-6]; CORT=7 [5.5-8]; Mann Whitney U test p=0.0014), reduced total grooming time (median grooming time [seconds] VEH=47 [IQR 26-67]; CORT=23 [7-42]; Mann-Whitney U Test p= 0.0120), and increased latency to feed in a novel arena indicative of higher anxiety (median latency to feed [sec] VEH=80 [IQR 42-194]; CORT=206 [162.5-315.5]; Mann-Whitney U Test p<0.001) (**Figure 1A-C**).

**Figure 1.**
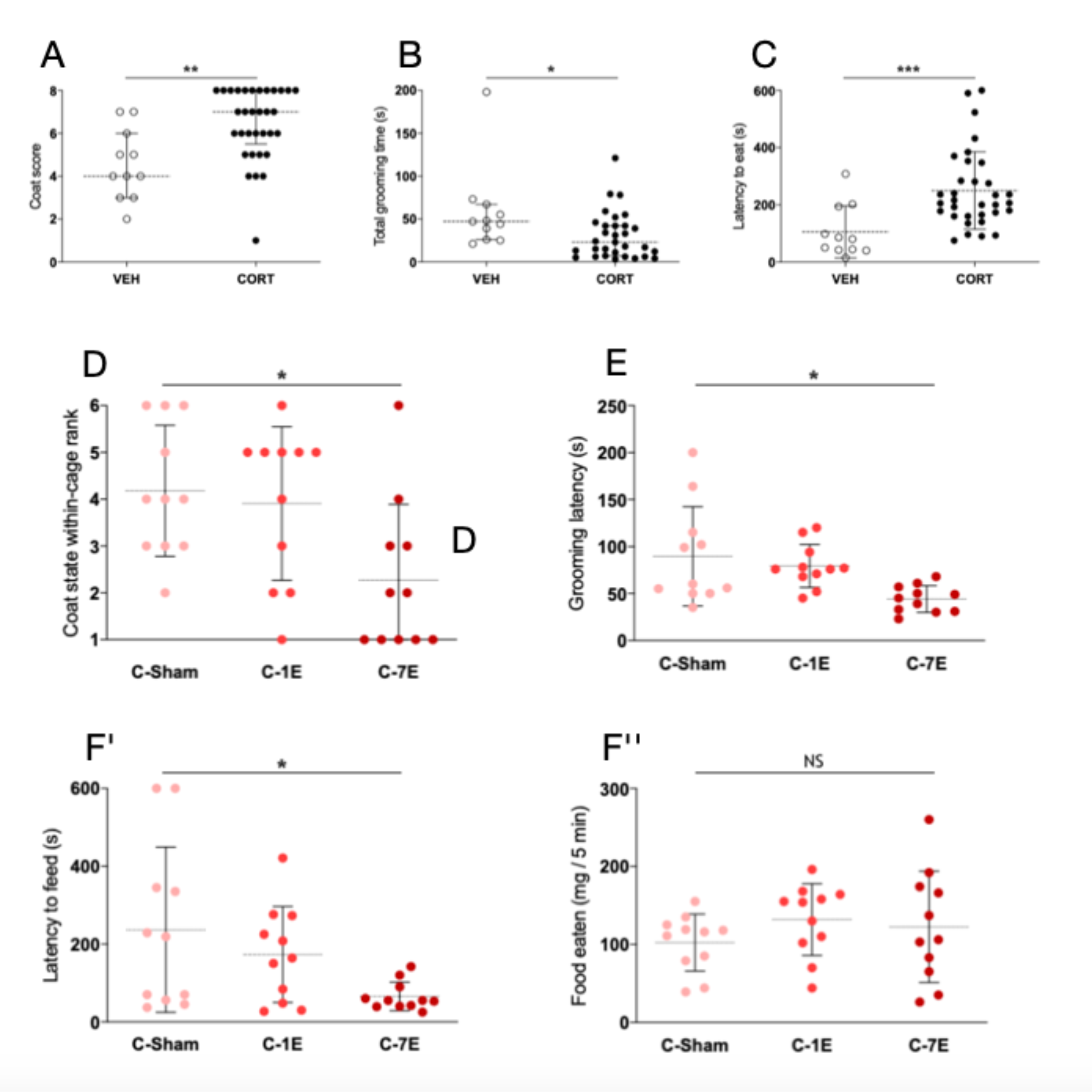
Repeated ECS reverses depressive-like behavior in a neuroendocrine model of depression. Note: **A-C** are behavioral data from T0 (CORT vs VEH-treated animals). **D-F** are behavioral data post-ECS (CORT animals only). **A:** Coat State test. CORT-treated animals had significantly poorer coat state. **B:** Splash Grooming test. CORT-treated animals showed a significantly lower total grooming time. **C:** Novelty-Suppressed Feeding. CORT-treated animals took significantly longer to eat in a novel anxiogenic environment. **D:** Effect of ECS on Coat State. Animals receiving 7ECS ranked significantly higher for coat state quality compared to cage-mates who had received 1ECS or Sham. **E:** Effect of ECS on Splash Grooming test. 7ECS-treated animals showed reduced latency to groom compared to 1ECS or Sham. **F’:** Effect of ECS on Novelty- Suppressed Feeding. Animals receiving 7ECS showed reduced time to eat in a novel anxiogenic environment, but no difference in total food eaten in their home cage (**F’’**), compared to 1ECS or Sham. N=11-33 animals per group. C-Sham= CORT then 7 sham stimuli. C-1E= CORT then 1ECS / 6 sham stimuli. C-7E= CORT then 7ECS. *p<0.05, **p<0.01; ***p<0.001. VEH= Vehicle treated animals. CORT= Corticosterone treated animals.

To identify a suitable ECS protocol that reverses this depressive-like state, we compared three ECS treatment groups; CORT plus 7 stimuli [CORT-7ECS], CORT plus one stimulus then six sham stimuli [CORT-1ECS], or CORT plus seven sham stimuli [CORT-Sham]. There was evidence of improved coat state only in animals receiving CORT-7ECS (median within-cage rank Sham=4th [IQR 3rd-6th]; 1ECS=5th [2nd-5th]; 7ECS=2nd [1st-3rd], Kruskal-Wallis Test p=0.020). Animals receiving 7ECS showed behavioral recovery of grooming (median latency to groom [sec] Sham=60 [IQR 50-115]; 1ECS=76 [68-94]; 7ECS=45 [31-57]; One-way ANOVA p=0.011) and feeding (median latency to eat [sec] Sham=219 [IQR 56-345]; 1ECS=164 [48-273]; 7ECS=55 [40-90]; One-way ANOVA p=0.0284) (**Figure 1D-F**). Therefore, consistent with previous studies (16), chronic CORT administration induced depressive-like behavioral deficits which were reversed by 7ECS.

### ECS preferentially induces neurogenesis in the ventral hippocampus in correlation with behavioral response

Adult hippocampal neurogenesis may be one mechanism underpinning antidepressant treatment effects. (3–5) We first confirmed, therefore, that ECS increased hippocampal neurogenesis in both CORT- and VEH-treated animals. We quantified the extent of ECS-induced neurogenesis by examining NeuN+EdU+ co-labelling in the dentate gyrus Granule Cell Layer (GCL) one month after treatment. In CORT-treated animals there was a highly significant effect of 7ECS (mean GCL NeuN+EdU+ density, VEH-treated ShamECS=2.8/mm3 [SD= 0.9]; CORT ShamECS=4.0/mm3 [1.4]; CORT-1ECS=4.9/mm3 [1.2]; CORT-7ECS=11.1/mm3 [1.7]; One-way ANOVA comparing CORT groups p< 0.0001) (**Figure 2D-E**).

**Figure 2.**
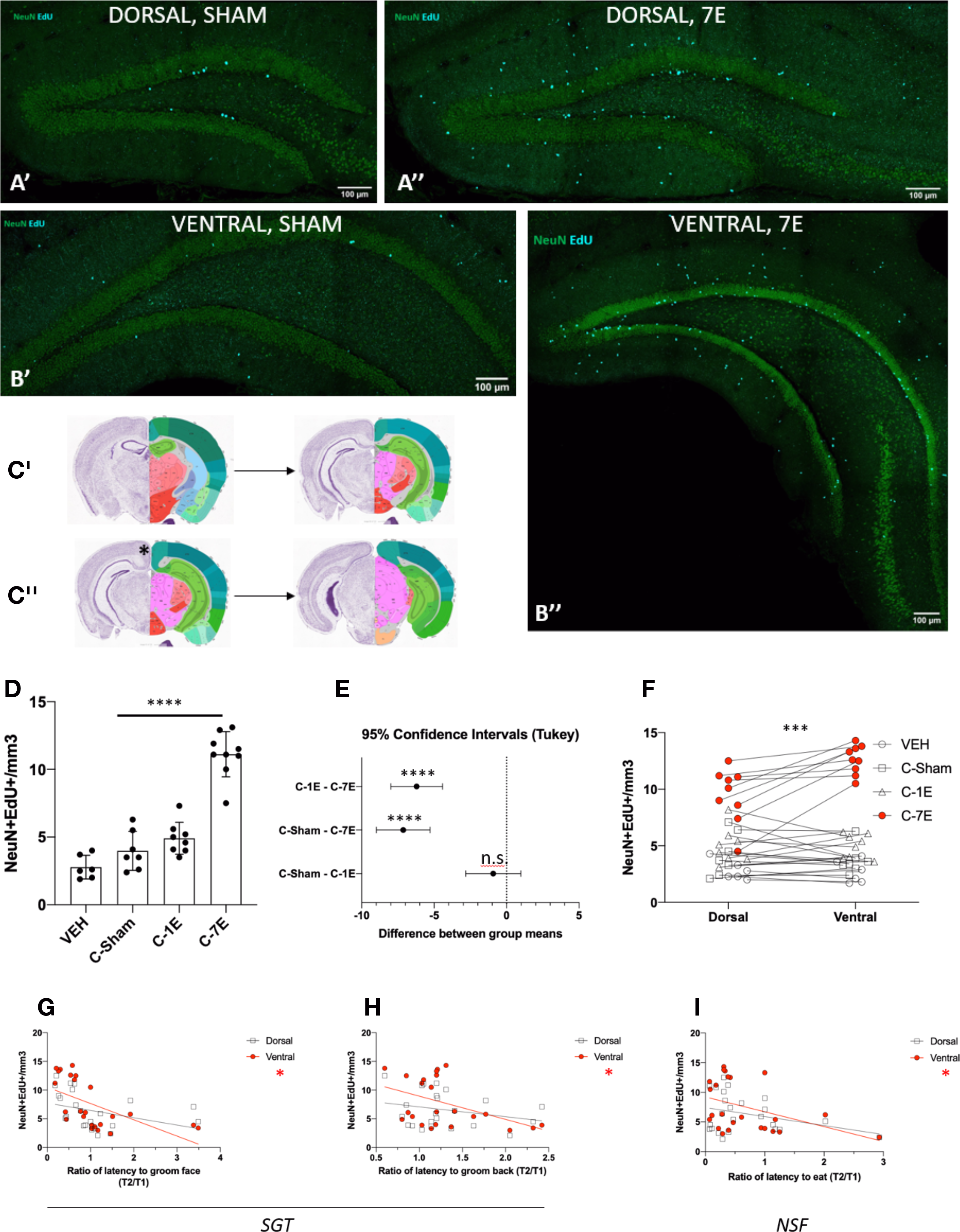
In depressive-state animals, repeated ECS induces neurogenesis preferentially in the ventral hippocampus in correlation with behavioral response. A: Representative images of newborn neurons (NeuN+EdU+ cells) in the dorsal dentate gyrus after Sham (A’) or 7ECS (A’’). **B:** Representative images from the ventral dentate gyrus after Sham (B’) or 7ECS (B’’). Note the qualitatively greater response of neurogenesis in the subgranular zone, of 7ECS over Sham in ventral as compared to dorsal sections. **C:** Schematic overview of dorsoventral allocation. Images are from the Allen Brain Atlas. The point at which the corpus callosum no longer crosses the midline (**\***) was used to separate dorsal (C’) from ventral (C’’) slices. **D:** Overall quantification of newborn neuronal density along the whole axis combined, in Sham,1ECS, or 7ECS groups. CORT- treated groups are compared using one-way ANOVA. **E:** Post-hoc Tukey’s test comparing differences in the density of neurogenesis between CORT-treated group pairs in (D). **F:** Density of ECS-induced neurogenesis in dorsal versus ventral compartments within the same animal. For VEH, CORT-Sham, and CORT-1E groups, densities between dorsal and ventral are roughly equal. By contrast CORT-7E animals had markedly increased density of neurogenesis in the ventral hippocampus. Tested using 2-way ANOVA. **G:** Correlation between ECS-triggered dorsal (black data points) and ventral (red data points) neurogenesis and Splash Grooming Test latency to groom face. **H:** Correlation between dorsoventral neurogenesis and Splash Grooming Test latency to groom back. **I:** Correlation with Novelty-Suppressed Feeding latency to eat. All analyses in **G-I** show significantly non-zero slopes for ventral but not dorsal neurogenesis. ***p<0.001; ****p<0.0001. N= 6-9 animals per group with at least eleven 30-micron sections analyzed per animal.

Animals receiving 7ECS showed increased density of neurogenesis throughout the length of the DG, with a greater increase in the ventral hippocampus (significant interaction between axis position and treatment group; 2-way ANOVA p<0.001; Sidak’s multiple comparisons test mean increase in 7ECS ventral hippocampus=3.2/mm3 [95%CI=1.6-4.7] relative to dorsal, p<0.0001) (**Figure 2A-C** and **2F**). A significant correlation was seen in the ventral hippocampus with the Splash Grooming test “latency to groom” face (linear regression, dorsal slope= -1.28 [95%CI -2.78 to +0.23], p=0.093; ventral slope = -2.84 [-4.60 to -1.07], p=0.003) and back (dorsal slope= -1.71 [- 4.59 to +1.17], p=0.231; ventral slope= -4.05 [-7.64 to -0.45], p=0.029), and with Novelty Suppressed Feeding (NSF) latency to eat (dorsal slope= -1.51 [-3.41 to +0.38], p=0.112; ventral slope= -2.50 [-4.98 to-0.03], p=0.048) (**Figure 2G-I**).

We concluded that in our model depressive state, multiple ECS were required to increase neurogenesis and reverse depressive-like behavior. Repeated ECS stimulated ventral hippocampal neurogenesis to a greater extent than dorsal, and this correlated with behavioral response. On the basis of these results, we only administered 7ECS (or 7Sham) when giving ECS in the remaining experiments.

### ECS impacts on a range of cell-types in the ventral SGZ

To examine which SGZ cell types responded to ECS, we delivered 7ECS or Sham to VEH-treated or CORT-treated animals. We performed RNA sequencing of ventral SGZ samples from these animals and two pre-ECS ‘baseline’ groups (n=4 animals per group) (**Fig S2**). Confirming the validity of this approach, exploratory Principal Components Analysis (PCA) revealed the strongest source of variance in the data to be the delivery of 7ECS. Naive and depressed-state ECS recipient animals also discernibly separated from each other while sham control animals in either stream did not (**Fig 3A-B**).

**Figure 3.**
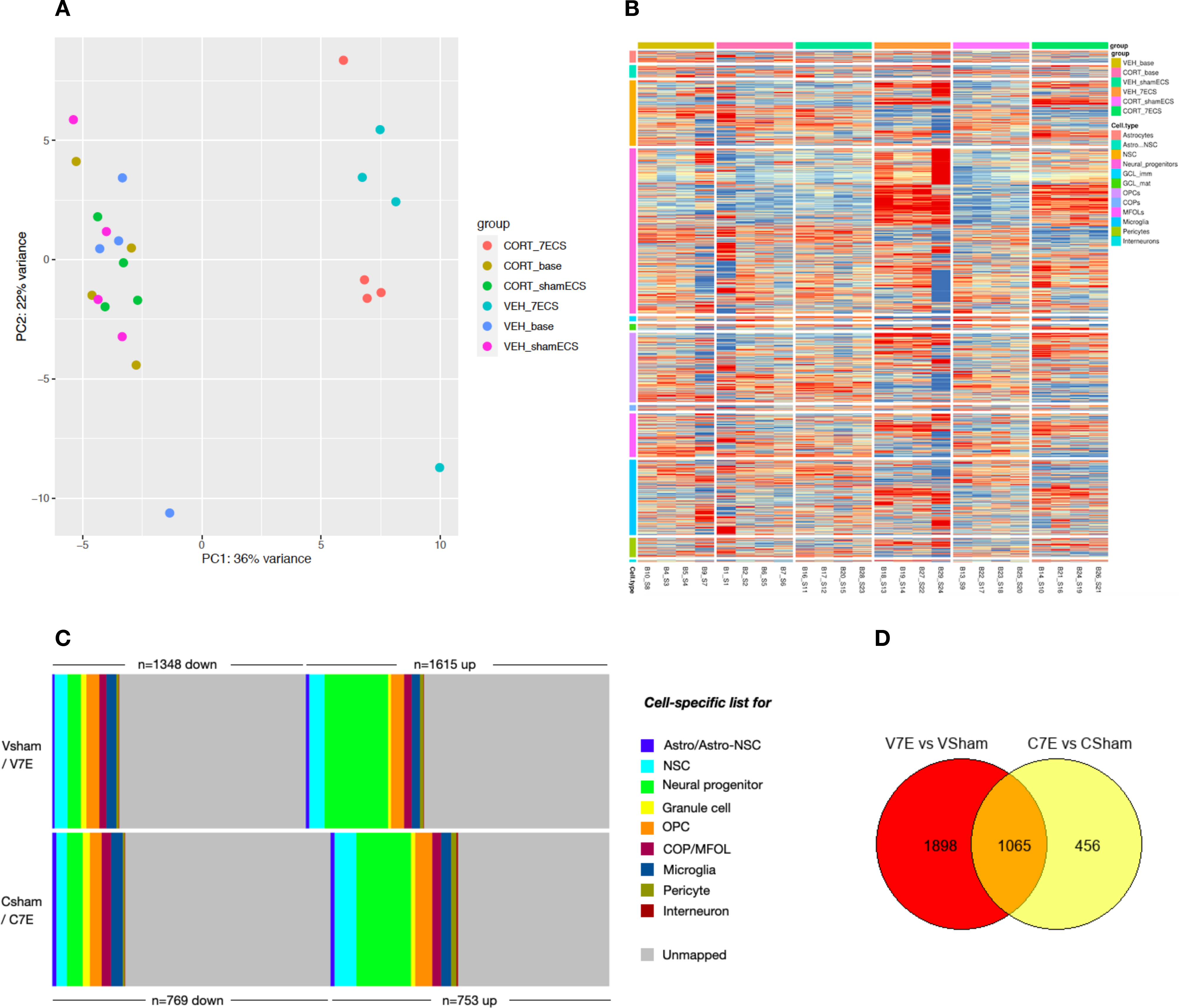
ECS preferentially affects NSPCs but also a range of cell-types in the ventral SGZ, with only partial overlap between naive and depressive-state animals. A: Principal Components Analysis of all six experimental groups (see **Fig S2**, VEH baseline, CORT baseline, VEH-7ECS, VEH-Sham, CORT-7ECS, CORT-Sham). ECS is the strongest signal of variance in the data. **B:** Heatmap of SGZ cell-type specific genes curated from single cell RNA-seq datasets (see **Table S1**). **C:** Stacked bar chart illustrating the proportion of up- and down-regulated genes after ECS in naive (VEH) and depressive-state (CORT) animals that can be mapped to cell-type specific gene lists. Note the striking similarity of proportions favoring up-regulated genes in NSPCs, but also extending across other cell types. For data see **Table S2**. **D:** Venn diagram illustrating only a partial intersect of ECS responsive DE genes in naive VEH-treated animals (V7E vs VSham) and depressive-state CORT-treated animals (C7E vs CSham).

We derived cell-type specific SGZ gene lists from literature (**Table S1** and Methods) and intersected these with lists of genes which were significantly up- and down-regulated after ECS. This comparison confirmed the impact of ECS on neural stem and progenitor cells in both VEH-treated and CORT-treated animals (**Fig 3C** and **Table S2** (note multiple tabs)). To test all cell types objectively, we constructed 2x2 tables comparing the proportion of genes for a given cell type which were differentially expressed in our dataset, against all differentially expressed genes as a proportion of the entire mouse genome. As shown in **Table S2**, Chi square Bonferroni-corrected p values were highly significant for NSC, NSPC, OPCs, microglia, committed oligodendrocyte progenitors/early myelinating oligodendrocytes, pericytes, and granule cells. In both VEH- and CORT-treated animals, ECS triggered both up- and down-regulation of cell-specific genes in nearly all these cell types. Astrocyte and Astrocyte-NSC gene lists showed weaker effects that did not pass Bonferroni correction. These data suggest that ECS has effects across a range of cell types in the ventral SGZ niche, not just neural progenitors, with a bias towards gene up-regulation rather than down-regulation.

### ECS promotes gene sets implicated in neural plasticity

In total, ECS induced the differential ventral SGZ expression of n=2,962 genes in VEH-treated (“naive”) animals, and of n=1,520 genes in CORT-treated (“depressive-state”) animals, versus respective Sham controls. Of these, n=1,064 genes were differentially regulated in both naive and depressive-state animals (**Fig 3D**). To identify enriched pathways, we subjected differentially expressed gene lists from naive and depressive-state animals to overrepresentation analysis. Results of these separate analyses are presented in **Table S3** (for naive animals) and **Table S4** (for depressive-state animals; both versus their appropriate Sham control). We then intersected these separate overrepresentation analyses with the aim of identifying specific Gene Ontology (GO) terms shared between VEH-treated and CORT-treated animals after ECS. We reasoned that such shared ECS-responsive pathways may be ‘core effects’ of the electrical stimulus and seizure, and therefore independent of prior depressive state (see Supplementary Methods).

The results of this analysis are presented in **Table S5**. Shared GO Biological Processes (n=60) included: metabolic or biosynthetic processes for cholesterol and glycolysis; axonogenesis; dendrite development; cell proliferation, synapse assembly, plasticity and transmission; cell-cell adhesion; and gene sets involved in learning, memory, and forebrain development. Shared ‘GO Molecular Function’ sets (n=15) included: semaphorin receptor binding (the top hit by Fold Change in both streams); cytoskeletal, actin, and extracellular matrix genes; and activity of voltage-gated ion channels. Shared ‘GO Cellular Component’ gene sets (n=28) included: myelin sheath (the top hit by FDR in both streams); pre-synaptic and post-synaptic membranes; synaptic vesicle and endocytic vesicles; neuron projection, and dendritic spines; extracellular matrix; adherens junction; and voltage-gated potassium channels. Because changes in these gene sets were seen after ECS in VEH-treated animals as well as CORT-treated animals, these results suggest that a significant component of the ventral SGZ response to ECS may be independent of disease state. ECS promotes highly metabolic cellular processes inherent to central nervous system plasticity, including cellular proliferation and migration, synapse assembly and transmission, extracellular matrix interactions, and growth factor signaling. Novel candidates for inclusion in these processes include semaphorin signaling, cell adhesion, and myelin gene sets.

### ECS reduces neuro-inflammation

We were interested to note that molecular pathways enriched by ECS only in depressive-state animals (as opposed to those shared between naive and depressive state animals) included cytokine/chemokine pathways; angiogenesis; and TGF-beta pathways (**Table S6**), suggesting a state-specific effect of ECS on neuro-inflammation. We therefore conducted Gene Set Enrichment Analyses (GSEA) comparing the SGZ sequenced output of CORT- with VEH-treated animals, and CORT-Sham with CORT-ECS animals. Unlike overrepresentation analysis, GSEA is not dependent on the pre-selection of differentially expressed genes, but instead utilizes data from every gene identified in the sequencing analysis output. Manually intersecting complementary GSEA output (see Methods) permitted us to discover SGZ gene sets perturbed by CORT then altered in the reverse direction by CORT-ECS.

As internal validation and as expected, after CORT (vs. VEH) we found a down-regulation of gene sets reflecting neurogenic metabolic activity (mitochondria, oxidative phosphorylation, and protein binding, processing, and transport gene sets) which were then up-regulated by CORT-ECS (vs CORT-Sham). We noticed that the top 20 gene sets (ranked by normalized effect size) up-regulated by CORT *and* the top 20 gene sets subsequently down-regulated by CORT-ECS included six immune response gene sets: GO BP “Immune Response”; Kegg “Cytokine-Cytokine Receptor Interaction”; Kegg “Complement and Coagulation Cascades”; Wiki-pathways “Complement and Coagulation Cascades”; TFACTS “NFKB1”; and GO MM “Response to Lipopolysaccharide”. These analyses suggest a dynamic and reciprocal regulation of neuro-inflammatory gene sets which are elevated by CORT - i.e., in depressive-state animals - and then reduced by CORT-ECS. The inflammatory / anti-inflammatory reciprocal molecular signal was apparent in both SGZ and PFC (**Table S7**).

We next searched for molecular signals of microglial activation status after ECS. We used microglial-specific genes differentially expressed after ECS (as determined by cross-referencing **Table S2** and the Barres sequencing database) as input to the Microglia Single-Cell Atlas of Health and Injury. (17) From a small list of five such genes (*Sgk1, Btg2, Cd83, Atf3, Ly6e*), we noted ECS-induced upregulation of the microglial activation inhibitor *Sgk1*, and down-regulation of the microglial activation marker *CD68* (18), in both CORT-7ECS and VEH-7ECS treated animals.

To further interrogate a role for ECS in reducing neuro-inflammation we used immunofluorescence to examine microglial staining patterns in the dentate gyrus of an independent cohort of animals treated with CORT and then 7ECS or Sham. After CORT, ECS reduced IBA1+ microglial density (mean IBA1+ density, CORT-Sham= 83.9/mm2 [SD= 6.0]; CORT-ECS= 67.3/mm2 [8.5]; n=4-5 animals per group, independent samples t-test p= 0.012). ECS also reduced the mean fluorescent intensity of IBA1+ individual cells (mean IBA1+ cell intensity, CORT-Sham= 82.9 [SD= 3.6]; CORT-ECS= 72.2 [6.5]; n=4-5 animals per group, independent samples t-test p= 0.022; n=4-5 animals per group where each animal’s data point represents the mean intensity of 27-75 individual cells), and returned the morphology of DG microglia to a ramified ‘resting state’, unlike the activated morphology seen in CORT-Sham treated animals (**Figure 4**).

**Figure 4.**
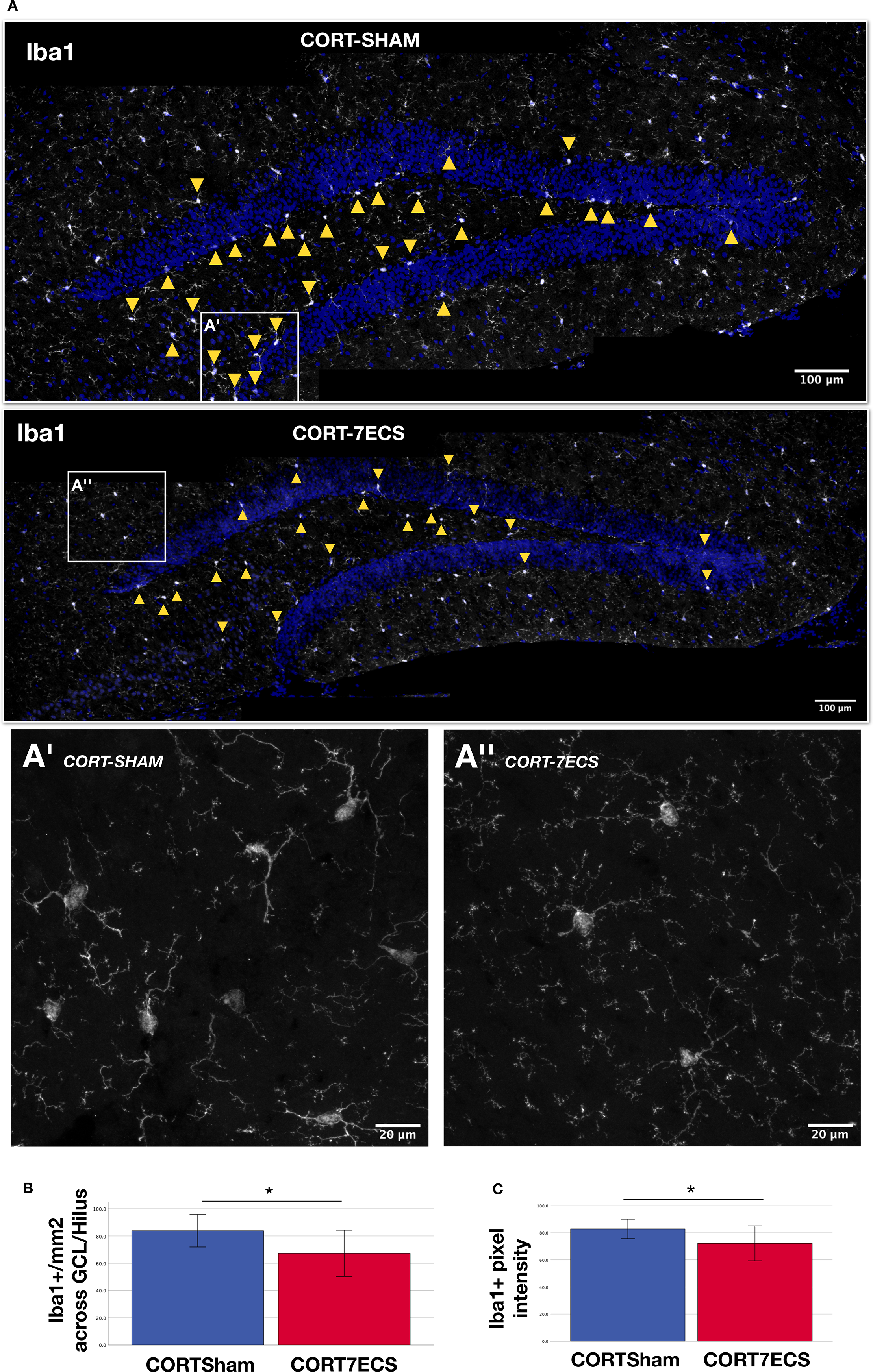
ECS reverses neuro-inflammation after chronic corticosterone. **A:** Immunofluorescence images illustrating microglial (Iba1+) density in the dentate gyrus (DG) in CORT-Sham and CORT-7ECS conditions. Arrowheads mark microglia in the GCL, SGZ, and Hilus. **A’** High magnification image of DG microglia in CORT-Sham treated animals, showing an ‘activated’ morphology. **A’’** Image of DG microglia in CORT-ECS treated animals, showing a more ‘resting’ morphology with ramified processes. **B:** Quantification of microglial density in the GCL, SGZ, and Hilus. 4-5 animals per group, 4-6 DG sections per animal. **C:** Quantification of microglial fluorescence intensity. 4-5 animals per group, 27-75 Iba1+ cells per animal.

To explore potential for human translatability, we cross-referenced ECS-regulated microglial genes (listed in **Table S2**) against genes known to be dysregulated in peripheral inflammatory cells of patients with Major Depressive Disorder. (19) This intersection revealed nine microglial genes dysregulated in depressive-state mice (*Asph, Cited2, Ier5, Smap2, Capn3, Cd83, Cebpa, Cyth4, Fmnl1, Golm1, Parvg, Tbxas1,* and *Zyx*) which are also differentially regulated in the immune system in depressed human patients. Five of these genes (*Cited2, Cd83, Cebpa, Cyth4, and Parvg*) have known roles in microglial activation (20–24); in mice, ECS modulated each in the direction of lessening activation status. In total therefore our data suggest that electroconvulsive stimuli reduce neuroinflammation and may reduce microglial activation.

### Identification of novel candidate “antidepressant-response” genes in the ventral SGZ

To identify additional candidate genes in the ventral SGZ that might mediate the recovery process from depression, we intersected three differential expression lists: baseline (CORT vs VEH), naive ECS (VEH-7ECS vs VEH-Sham) and depressive-state (CORT-ECS vs CORT-Sham) (**Fig 5A**).

**Figure 5.**
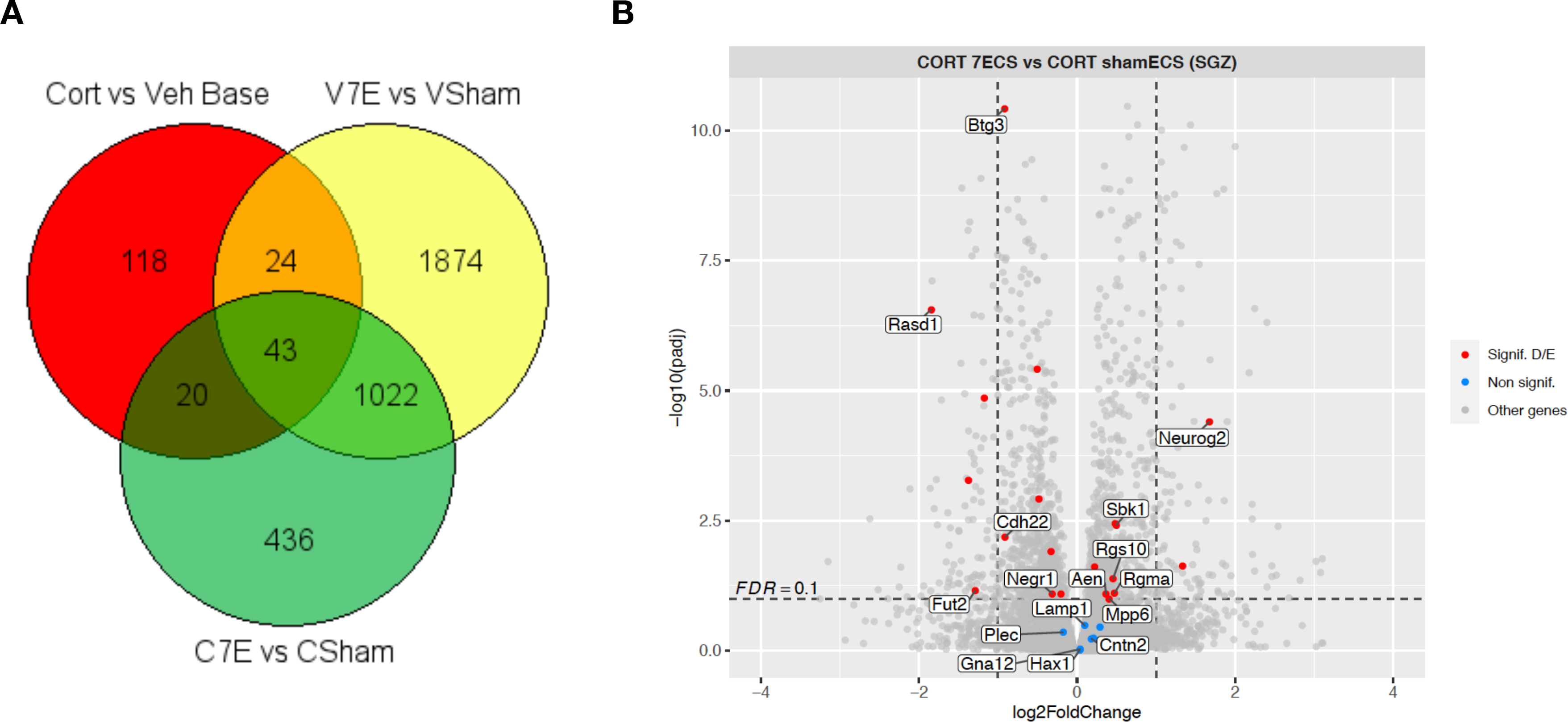
Identification of candidate “antidepressant-response” genes in the ventral SGZ. A: Venn diagram illustrating the intersect of genes dysregulated by CORT (CORT vs VEH base), and also differentially expressed after ECS in naive VEH-treated animals (V7E vs VSham) and depressive-state CORT-treated animals (C7E vs CSham). Genes falling within the overlap between CORT baseline and all other groups are listed under the appropriate headings in Table S7 (n=86 genes after excluding one label common to all analyses, ‘NA’). Note that some genes present in the overlapping areas of this diagram were differentially expressed in the same direction, and others in reciprocal directions. **B:** Volcano plot highlighting genes that were dysregulated by CORT and reversed by depressive-state ECS and/or by naive state ECS. Within this list, only those associated with neuropsychiatric outcomes in two or more human genomic databases are labelled with gene name. The plot presented is for the CSham vs C7E comparison. The VEH vs CORT and VSham vs V7E comparisons, with different values for each gene, are presented in Figure S3. Red points indicate significantly differentially regulated genes in the comparison shown. Some genes may only be significant in one or other of the ECS streams: where this is the case (e.g., if they were significantly regulated only after V7E), they are colored blue if non-significant in (e.g.) the plot for CSham vs C7E.

Selecting only those genes that were differentially regulated by CORT and specifically reversed by ECS after CORT revealed n=44 candidates of particular interest as a potential “antidepressant footprint”. A further 12 CORT-dysregulated genes were reversed by ECS in VEH-treated animals only; to maximize sensitivity to detect ECS-responsive genes, we included them in the present analysis. (**Table S8**).

Some of these genes have known functional roles in hippocampal neurogenesis (e.g., *Neurog2, Sox11, Tubb3,* and *Calb2,* all reduced by CORT and increased by ECS) and may simply be consequent upon the reduction and recovery of NSPCs in the experimental model. Others are not known to have a role in neurogenesis, but present biologically plausible candidacies including: *Cryab* (which suppresses neuro-inflammation, (25) reduced by CORT and restored by depressive-state ECS only); *Lynx1* (a negative regulator of synaptic plasticity and cognition, (26) elevated by CORT and reduced by depressive-state ECS); and *Slc29a1* (expressed in neural progenitors (**Table S1**) and a negative regulator of proliferation, (27) elevated by CORT and reduced by depressive-state ECS).

Finally, to explore the relevance of these 56 candidate ‘antidepressant response’ genes to human neuropsychiatric disorder, we intersected them with three human genomic databases: the EMBL-EBI GWAS catalogue; (28) the FUMA GWAS platform; (29) and an Epigenome-Wide Association Study (EWAS) Atlas. (30) Notably, we found that half (n=28) have been linked to human neuropsychiatric phenotypes in genome- or epigenome-wide association studies. These phenotypes include: sleep (*Hax1, Lamp1, Aen, Rasd1*); cognition (*Negr1, Sbk1, Cntn2, Cdh22, Ckap2l, Gap43*); inflammation (*Slc29a1*); white matter integrity (*Trabd2b, Btg3, Gna12, Plec*); response to psychiatric treatments (*Ckap2l, Gap43*); schizophrenia (*Mpp6, Cdh22, Hax1, Cntn2*); ASD (*Mpp6, Lamp1, Fut2*); bipolar affective disorder (*Hax1, Lamp1, Plec, Fut2*) and depressive symptoms (*Negr1, Rgs10, Rgma, Lynx1*) (**Fig 5B**, **Table 1**). They represent novel candidate molecular mechanisms that may underpin recovery from depression, and which warrant future study.

**Table 1.** Intersecting candidate genes with human GWAS and EWAS databases. *This Table describes 40 genes which were:* *1) differentially regulated by CORT ve VEH (FDR<0.05), **and*** *2) differentially regulated by ECS vs SHAM (after VEH or CORT, FDR<0.05; n=82, see Fig 5a), **and also*** *3) called within a FUMA gene-set analysis for neuropsychiatric outcomes, **or*** *4) called individually in the EBI-EMBL GWAS catalogue,* ***or*** *5) called individually in the Chinese National Genomics Data Centre EWAS Atlas*. *To determine FUMA gene-sets, full DGE results from V7E and C7E were run separately through the FUMA platform. Neuropsychiatric gene sets called in the results were extracted and intersected individually with the list of 82 potential candidate genes identified via Fig 5a*.

| Cort | V7E | C7E | FUMA | EBI | EWAS | Gene | Associated in human studies |
| --- | --- | --- | --- | --- | --- | --- | --- |
| Down | Up | - | o | o | o | Hax1 | BPAD, SCZ, sleep phenotype |
| Down | Up | - | o | o | o | Lamp1 | BPAD, Alzheimer's, sleep phenotype, ASD |
| Up | Down | Down | o | o | - | Negr1 | MDD, cognition |
| Down | Up | Up | o | o | - | Sbk1 | Cognition |
| Up | Down | Down | o | o | - | Rasd1 | Chronotype |
| Down | - | Up | o | o | - | Mpp6 | Mixed neuropsychiatric, SCZ, neuroticism, cognition |
| Up | Down | - | o | o | - | Plec | BPAD, white matter microstructure |
| Down | Up | - | o | o | - | Cntn2 | SCZ, cognition |
| Down | Up | Up | o | - | o | Neurog2 | ALS, SCZ |
| Up | Down | Down | o | - | o | Fut2 | BPAD, ASD |
| Down | - | Up | - | o | o | Aen | Chronotype (sleep duration), suicidal behaviour |
| Up | - | Down | - | o | o | Cdh22 | Cognition, SCZ endophenotype, neurodevelopment |
| Down | - | Up | - | o | o | Rgma | Chronotype, BPAD/MDD |
| Down | Up | Up | - | o | o | Rgs10 | Depressive symptoms, neurodevelopmental disorder |
| Down | Up | - | - | o | o | Gna12 | Brain volume, WM integrity, neurodevelopmental disorder |
| Up | Down | Down | - | o | o | Btg3 | White matter volume, maternal depression |
| Up | - | Down | o | - | - | Etnppl | ALS |
| Down | Up | Up | - | o | - | Ckap21 | Cognition (working memory) |
| Down | Up | Up | - | o | - | Gap43 | BPD, anxiety (response to treatment), cognition |
| Up | - | Down | - | o | - | Slc29a1 | Inflammation (Interleukin-10) |
| Up | Down | Down | - | o | - | Trabd2b | White matter integrity, reaction time |
| Down | Up | - | - | o | - | Abcb10 | Cortical volume |
| Down | Up | - | - | o | - | Bcas1 | BPAD, SCZ, ADHD |
| Down | Up | Up | - | - | o | Ptges3 | Neurodevelopmental disorder |
| Up | - | Down | - | - | o | Lynx1 | MDD |
| Up | - | Down | - | - | o | Rnf31 | SCZ |
| Up | Down | Down | - | - | o | Ebf4 | Neurodevelopmental disorder |
| Up | Down | Down | - | - | o | Prr36 | Neurodevelopmental disorder |

**Table 1. Intersecting candidate genes with human GWAS and EWAS databases.**
| Cort | V7E | C7E | FUMA | EBI | EWAS | Gene | Associated with (in humans) |
| --- | --- | --- | --- | --- | --- | --- | --- |
| Up | - | Up | o | o | o | Nr2f2 | Chronotype, alcoholism endophenotype, SCZ |
| Down | Down | - | o | o | o | Yipf4 | Chronotype, cognition, Alzheimer's |
| Down | Down | - | o | o | o | Prr16 | Depressed mood, cognition |
| Up | Up | Up | o | o | - | Ldlr | MDD, sleep duration, cortisol measurement |
| Down | Down | Down | o | o | - | Rasd2 | Chronotype, cognition |
| Up | Up | - | o | o | - | Itih3 | BPAD, SCZ, PD |
| Down | Down | - | o | o | - | Prdm5 | Cortical volume |
| Down | Down | Down | - | o | o | Spata13 | Cerebral growth, attention, depression, Alzheimer's |
| Down | Down | - | - | o | o | Rfx2 | BPAD, cognition, Alzheimer's |
| Up | Up | Up | - | o | - | Eya1 | Autism, sleep phenotype |
| Up | Up | - | - | o | - | Cdhr3 | Alzheimer's |
| Down | Down | - | - | o | - | Plekhh1 | Kynurenine levels in MDD |

## DISCUSSION

Despite being a fast, safe, and effective treatment for severe depressive illness, ECT has a negative public image. Alongside misleading depictions in the media and film/TV industries, stigma may also persist because mechanistic explanations remain elusive. Here we presented experiments suggesting that the behavioral response to repeated electroconvulsive stimuli correlates with adult neurogenesis most strongly in the ventral hippocampus. Many ventral SGZ molecular pathways induced by repeated ECS were shared between naive and depressive-state conditions while others were unique to the depressed state. Comparison of selectively enriched pathways suggested that ECS reduces neuro-inflammation. We further identified a novel list of candidate ‘antidepressant response’ genes, of which a high proportion have been implicated in human neuropsychiatric phenotypes.

Repeated ECS stimulated neurogenesis throughout the dorsoventral axis of the hippocampus but most strikingly, in depressive-state animals, in the ventral hippocampus. This observation correlated with behavioral recovery and is consistent with prior work suggesting functional dissociation along the hippocampal dorsoventral axis. (31) Cognitive functions may localize dorsally, and antidepressant-related effects more ventrally. (32) Since ECT treats depressive symptoms but may also cause episodic memory deficits, (33) the question of its molecular effects along the entire hippocampal axis deserves future study.

Molecular analysis of the ventral SGZ response to ECS revealed a striking number of ECS- responsive genes and pathways shared between naive and depressive-state animals. These shared pathways were enriched in multiple aspects of brain plasticity. Some (such as neurogenesis and synaptic plasticity) are previously well reported, while others (such as semaphorin signaling and myelination) are novel. Other pathways were relatively more enriched after ECS in the depressive-state, such as neuro-inflammatory and angiogenic pathways. This nuanced picture is a resource for hypothesis generation and highlights the value of disease modeling when studying antidepressant mechanisms.

Chronic stress increases microglial activation (34), and neuro-inflammation has been proposed to mediate the antidepressant response to ECT. (35) In keeping, we showed that ECS reversed molecular signatures of CORT-induced neuro-inflammation and led to a decrease in microglial density, staining intensity, and activated morphology in depressive-state animals. Our data are consistent with reports that ECS can alter microglial purinergic currents (36) and attenuate neuro- inflammation in a mouse model of multiple sclerosis. (37) Preclinical data on neuro-inflammation after ECS remain sparse, however, particularly in studies that first model a depressive state (5), while a limited number of human studies so far conflict on whether serum inflammatory markers associate with response after ECT. (38, 39)

We also identified a novel set of 56 candidate genes in the ventral hippocampal SGZ whose expression levels changed with induction of a depressive-like state and were reversed with ECS. In particular eleven genes - *Sbk*, *Rasd1, Negr1*, *Mpp6, Neurog2, Fut2, Aen, Cdh22, Rgma, Rgs10,* and *Btg3* - were associated with neuropsychiatric symptoms in at least two of three large-scale human databases and were noted to ‘recover’ after ECS in depressive-state animals.

Although discussion of this entire list is out of scope, we wish to briefly highlight two such genes. *Rasd1* encodes a GTPase that is up regulated in the brain by stress (40) and negatively regulates cell proliferation and survival *in vivo*. (41,42) Our observation that *Rasd1* is induced by CORT and repressed by ECS now implicates this gene in the antidepressant response. *Sbk1* meanwhile encodes a protein kinase with a functional role in embryonic mammalian brain development. (43) Despite scattered indications of a neurotrophic role for *Sbk1*, (44) its function in the mammalian CNS remains incompletely understood. Intriguingly however its Drosophila homologue *Meng-Po* promotes CREB signaling to facilitate the retrieval of previously learned memories. (45) Our finding that *Sbk1* was reduced by CORT and restored by ECS suggests that *Sbk1* could have a role in mediating the cognitive sequelae of depression or its treatment.

Together, our data 1) suggest a preferential response of the ventral hippocampal SGZ to antidepressant treatments, 2) characterize ventral SGZ molecular pathways in naive and depressive-state conditions, 3) highlight a potential role for neuro-inflammatory pathways in mediating the antidepressant response to ECS, and 4) identify novel ‘antidepressant response’ genes implicated in human neuropsychiatric phenotypes. Future functional and behavioral experiments will unpick which state- and ECS-responsive genes or gene pathways contribute causatively to the antidepressant response, and which are correlative or reflect compensatory changes.

## Supporting information

Figure S1. Experimental design for behavioural experiments.

Figure S2. Experimental design for RNA-seq.

Table S1. Gene lists for SGZ heatmap (all cell types).

Table S2. Cell-specific gene lists intersected with DGE.

Table S3. VEH-Sham vs VEH-7ECS G.O output.

Table S4. CORT-Sham vs Cort-7ECS G.O. output.

Table S5. G.O. sets induced by 7ECS - shared.

Table S6. G.O. sets induced by 7ECS - CORT only.

Table S7. GSEA of reciprocally regulated gene sets.

Table S8. CORT regulated genes that respond to ECS.

## ACKNOWLEDGEMENTS

AGR was funded by a Royal College of Physicians of Edinburgh JMAS Sim Fellowship, an MRF/MRC PsySTAR Fellowship, and a University of Edinburgh / Wellcome Trust ISSF-3 consumables grant. This work has made use of the resources provided by the Edinburgh Compute and Data Facility (ECDF). We are grateful for the generous assistance of Dr. Caroline Stewart in enabling our model of ECS, animal unit, veterinary, imaging facility, and histology technical staff in the Centre for Regenerative Medicine, and members of the ffrench-Constant, Williams, and Miron laboratories for many helpful discussions.

## DISCLOSURES

The authors have no conflicts of interest to declare.

---

Figure S1. **Experimental design (behavioral and cellular analyses).** Eight-week-old animals received Corticosterone (CORT) or Vehicle (VEH) in the drinking water as detailed in Methods. Between D21 and D28, baseline behavioral tests (Coat State, Splash Grooming test, and Novelty Suppressed Feeding test) were conducted comparing CORT-treated animals with VEH-treated animals (T0). Starting on D29, all animals then received an ECS schedule every 48h for 14 days (7 treatments total). EdU was also injected every 48h during this time. From D43, the same behavioral tests were repeated on the CORT-treated groups while the animals continued CORT (T1). On D50 CORT was stopped and the animals were tested again in the following week (T2). All animals were followed up until one month following the last ECS treatment then sacrificed.

Figure S2. **Experimental design (molecular analyses).** Eight-week-old animals received Corticosterone (CORT) or Vehicle (VEH) in the drinking water as detailed in Methods. After five weeks both CORT and VEH treated streams were randomized to receive 7-ECS or 7-Sham, while CORT (or VEH) continued. Animals (n=4 per group) were sacrificed as detailed in Methods, before ECS began (for CORT baseline versus VEH baseline analysis), and two days after the final ECS or Sham (for VEH-7ECS versus VEH-Sham, and CORT-7ECS versus CORT-Sham analyses). Brains were processed for ventral SGZ RNA sequencing as detailed.

Figure S3. **Volcano plots for VEH/CORT, and VSham/V7E comparisons.** The corresponding VEH vs CORT and VSham vs V7E comparisons for the volcano plot analysis shown in Fig 5B. See Fig 5B for orientation.

TABLE S1. Cell-type specific genes from single-cell RNA-seq datasets of the SGZ niche.

TABLE S2. Cell type specific gene lists intersected with DGE results after ECS.

TABLE S3. VEH-7ECS vs Sham ECS (all DGE) unfiltered Gene Ontology output.

TABLE S4. CORT-7ECS vs Sham ECS (all DGE) unfiltered Gene Ontology output.

TABLE S5. G.O. sets induced by 7ECS in both naive (VEH) and depressive (CORT) states.

TABLE S6. G.O. sets induced by 7ECS in the depressive (CORT) state only.

TABLE S7. GSEA comparing reciprocally regulated SGZ gene sets.

TABLE S8. CORT dysregulated genes that also respond to ECS.

## Notes

### Competing Interest Statement

The authors have declared no competing interest.

