## Supplemental Methods for "Electroconvulsive stimuli reverse neuro-inflammation and behavioral deficits in a mouse model of depression"

*Corticosterone administration* CORT was dissolved by shaking for 4h at room temperature with the inert carrier vehicle 0.05% 2-Hydroxypropyl beta-cyclodextrin (Cayman Chemical 16169). Bottles were protected from light, changed every 2-3 days, and continued during ECS.

*Random allocation* Animals were randomly allocated to one of three treatment groups (CORT plus 7 stimuli [CORT-7ECS], CORT plus one stimulus then six sham stimuli [CORT-1ECS], or CORT plus seven sham stimuli [CORT-Sham]. Animals were randomised to these groups within cages to minimise cage effects.

*Coat State Test* Mice were examined by a blinded and experienced animal technician who scored each animal according to how unkempt their coat was. For baseline (T0) analyses we modified a scoring system that has been used by other groups. (1) Points were awarded on a 0-1-2 ordinal scale for an unkempt coat on each of the neck, back, right flank, and left flank areas, for a total possible score of 8 points. For longitudinal (T1 and T2) analyses however the same blinded technician reported feeling unable to reproduce the same baseline level of internal ‘calibration’ or rigour of inspection over time. For these later assessments therefore we took advantage of animals being randomised to ECS groups within each cage, to pilot a simpler ranking system whereby after the final ECS, animals in each cage were simply ranked from 1 (best coat) to 6 (worst coat) on the basis of the blinded technician’s experienced opinion. The spread of ranking scores between groups was analysed. This system had good face validity and was easy to implement.

*Novelty Suppressed Feeding* Mice were denied access to food for 16h prior to testing. They were then acclimatised to a quiet testing room and temporary holding cages for 1hr before testing. Animals were individually placed in an open field (60cm x 60cm) under bright lighting, with a piece of standard chow placed on a card in the centre of the field. The time taken to grasp the food, bite it and settle to eat, was recorded up to a maximum time of 600 seconds. The field was cleaned with 70% alcohol before each mouse was tested. To increase motivation animals were not allowed access to food for 16h before testing. To minimise cage effects caused by the passage of time and potentially longer fasting, one animal was tested per cage in cage series, repeating until all animals had been tested. The field was cleaned with 70% ethanol between each animal. The test was video-recorded for later corroboration of timings. Novelty was maintained during repeat testing by varying the shape and colour of the card in the centre of the field.

*Splash Grooming Test* This is a test of induced grooming following a splash stimulus. Initial trials of squirting water or sucrose solution on the animal’s coat using a 200 microlitre pipette were difficult to reproduce consistently, because the unkempt coat state induced by Corticosterone generally repelled water. Trials of lightly misting the animal’s snout with a single fine spray of water were also highly variable, with minor differences in the animal’s head position during the spray observed to have significant effects on grooming behaviour. However the passive drag of a saturated piece of towelling was highly reproducible. Towelling was cut to identical sizes, wettened with 10ml of autoclaved drinking water, and moved from side to side across the animal’s back six times under the influence of gravity only. The mouse was held gently by the tail on top of the cage and its instinctual reach and grab on the cage with its forepaws naturally exposed its back to the towelling. All animals tolerated this procedure well. The dampened mouse was placed in a tall glass cylinder backed by mirror panels to allow clear behavioural observations regardless of the animal’s orientation. Time to groom face, time to groom back, and total grooming time were analysed up to a maximum of 6 minutes.

*Tissue sectioning* Floating sections were cut from throughout the length of the dorsoventral axis of the dentate gyrus (Bregma -1.22 to -3.88) in a 1:6 ratio and 80 micron thickness. Laterality when floating was maintained by piercing the right hemispheric cortex with a fine 30G needle before sectioning. Sections were washed in 1xPBS for immediate use and stored long-term in 1xPBS with 0.01% azide at 4C.

*Confocal Microscopy* Images were taken with a Leica SP8 confocal microscope at 20X magnification in 30 micron stacks of 2 micron optical sections. Sequential scanning of channels was performed to avoid spectral bleed-through. Images were processed using FIJI freeware (<https://fiji.sc/>) and analysed blind to experimental group.

*Dorsoventral allocation* Sections were reconciled in a blinded manner to dorsoventral order by inspection of their gross morphology under a brightfield light microscope. A Mouse Brain Atlas (2) was consulted to interpret anatomical cues such as whether the corpus callosum was still present in the midline, the morphology of the cingulum and forceps major if not, the size and shape of the dentate gyrus granule cell layer, and the distribution of ventricular sulci. This process allowed reliable allocation of each section back to the reference atlas. In the Atlas the corpus callosum divides at between -2.54 and -2.70 relative to Bregma, a position half-way between the dorsal (-1.22) and ventral (-3.88) limits of our sectioning. Taking advantage of this equal split, sections with an intact corpus callosum were allocated to dorsal hippocampus and those where it had divided were considered to be ventral.

*Laser-capture microscopy* For SGZ sectioning, snap-frozen posterior brain blocks were mounted on their anterior face to allow a posterior approach and brought up to -12C in a Leica Cryostat (chamber -20C, head -12C). Eight micron coronal sections of the ventral hippocampus (Bregma -3.68 to -2.88) were cut to RNAse-free PEN slides (Zeiss) in a 1:10 series with six sections (twelve SGZ) per slide. Sections were firmly mounted by fingertip thawing, immediately re-frozen and kept on dry ice in closed 50ml Eppendorf tubes. At laser capture, slides were individually processed in series. Each slide was thawed by immersion in room temperature 70% RNAse-free ethanol (Sigma E7023) with RNAse-free water (Invitrogen 10977-035) and 0.5% Cresyl Violet (Acros Organics 229630050) for ten seconds, then 90% ethanol for 60 seconds, and 100% ethanol for 60 seconds. The slide was allowed to air dry at room temperature for three minutes and processed using a PALM laser capture microdissection microscope (Zeiss) on x20 magnification. The SGZ was defined as two granule cell nuclei widths above and below the hilar border. To maintain RIN quality the maximum time time to process each slide was limited to 25 minutes. This time frame allowed 24-30 SGZ to be dissected and collected in 200µl Adhesivecaps (Zeiss 415190-9191-000, one cap per slide) which were kept on dry ice until RNA extraction. For PFC sectioning, anterior brain blocks were mounted by the anterior pole to allow a posterior approach to PFC, and sectioned in eight micron coronal sections at -12C. Slides were processed for laser capture microdissection as above. Because the PFC is a much larger area than SGZ, four brain sections (eight PFC, Bregma +1.70-+2.00) were dissected for RNA extraction per animal.

*RNA quality and concentration* RNA quality and concentration were checked on an Agilent Tapestation 2200 using the High-sensitivity Screentape (Agilent 5067-5579). All SGZ samples sent for sequencing had RIN ≥ 7.5 (range 7.5-8.7, mean 8.3). Sample concentrations eluted in 16 microlitres of RNAse-free water ranged from 1.2ng/microlitre to 3.6ng/microlitre. All PFC samples had RIN ≥ 7.7 (range 7.7-8.8, mean 8.2, concentrations 0.9-3.3ng/microlitre).

*Data preprocessing* Read quality from individual RNA-seq libraries was assessed using FastQC (v.0.11.4) (3) and MultiQC (v.1.3.dev0). (4) Reads were pseudoaligned to the Mus musculus transcriptome (Ensembl version GRCm38.96; April 2019) using kallisto quant (Linux v.0.44.0). (5) Data was imported into R (v.3.6.3). A read count matrix for each area was computed using the tximport Bioconductor package (v.1.14.2). (6)

*RNA-seq analytic design* For SGZ RNA and PFC RNA separately, differential expression was computed for the following three pairwise comparisons: (i) VEH baseline vs CORT baseline; (ii) VEH-Sham ECS vs VEH-7ECS; (iii) CORT-Sham ECS vs CORT-7ECS.

*Pathway analysis pipeline* We developed a pipeline to interrogate ECS-induced pathways in VEH-treated and CORT-treated animals. First, lists of differentially-expressed genes (FDR<0.1 in pairwise comparisons) were added to the PANTHER classification system (7) to identify Gene Ontology (GO) biological processes, molecular functions, cell components, pathways, protein classes, and Reactome pathways overrepresented among the subset of significantly differentially expressed genes. Fold enrichment was computed by Fisher’s Exact Test; GO processes with FDR<0.05 were considered to be significantly enriched. Full hierarchical results were exported to tables. To avoid spurious enrichment results for gene sets with extreme numbers of genes, gene sets with <15 or >500 genes were excluded. The VEH-ECS/VEH-Sham and CORT-ECS/CORT-Sham outputs were then intersected; these intersected lists were manually curated to categorise each specific gene set either as “shared” between VEH and CORT after ECS, or “unique” to one. Because gene sets were not infrequently called within the same hierarchy (see e.g., “Sterol biosynthetic process” and “Cholesterol biosynthetic process”), the lists of shared and unique genes were then mapped back onto the full hierarchical results in order both to curate the most specific sole representative of each hierarchical branch, and to identify hierarchies unique to either VEH or CORT. This pipeline allowed us to summarise key shared and unique gene sets while minimising duplication or overlap as far as possible.

For each pairwise comparison in SGZ and PFC, we also interrogated the entire output ranked by log2 fold-change, using the Gene Set Enrichment Analysis platform. (8,9) This resulted in 12 GSEA output tables: up-regulated or down-regulated (2) in SGZ or PFC (x2) for each pairwise comparison (x3) = 12. Gene sets from these 12 outputs called at FDR <0.2 were tabulated. The tables were then manually intersected to identify gene sets meeting the following criteria: disordered by CORT (vs. VEH) **and** reciprocally reversed by CORT-ECS (vs. CORT-Sham), plus or minus similar effect in VEH-ECS (vs VEH-Sham). Lists of these reciprocally regulated gene sets were then constructed for those called only in the SGZ, **or** only in the PFC, **or** shared between both areas. This analysis allowed us to identify brain-wide, or region-specific, gene sets which were disordered in the model depressed state and then reversed by effective antidepressant treatment.
