## Supplementary figures and images for "Electroconvulsive stimuli reverse neuro-inflammation and behavioral deficits in a mouse model of depression"

### Figure S1. Experimental design for behavioural experiments.

Figure S1. Experimental design for behavioural / cellular analyses.

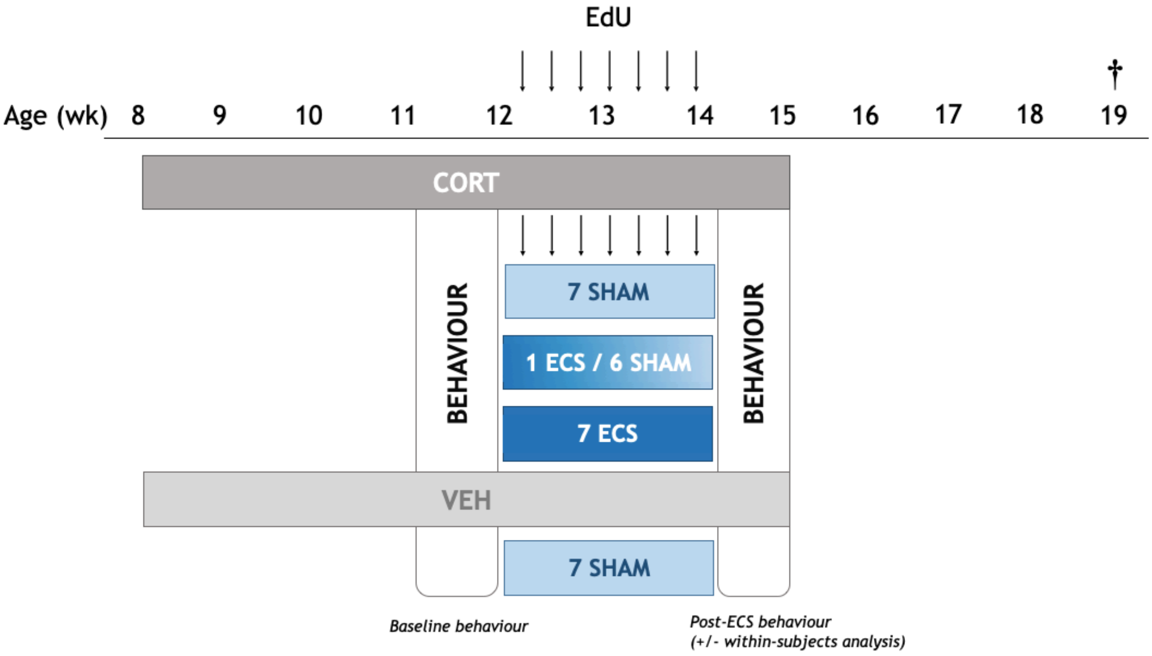

### Figure S2. Experimental design for RNA-seq.

FIGURE S2. Experimental design for molecular analyses (RNAseq)

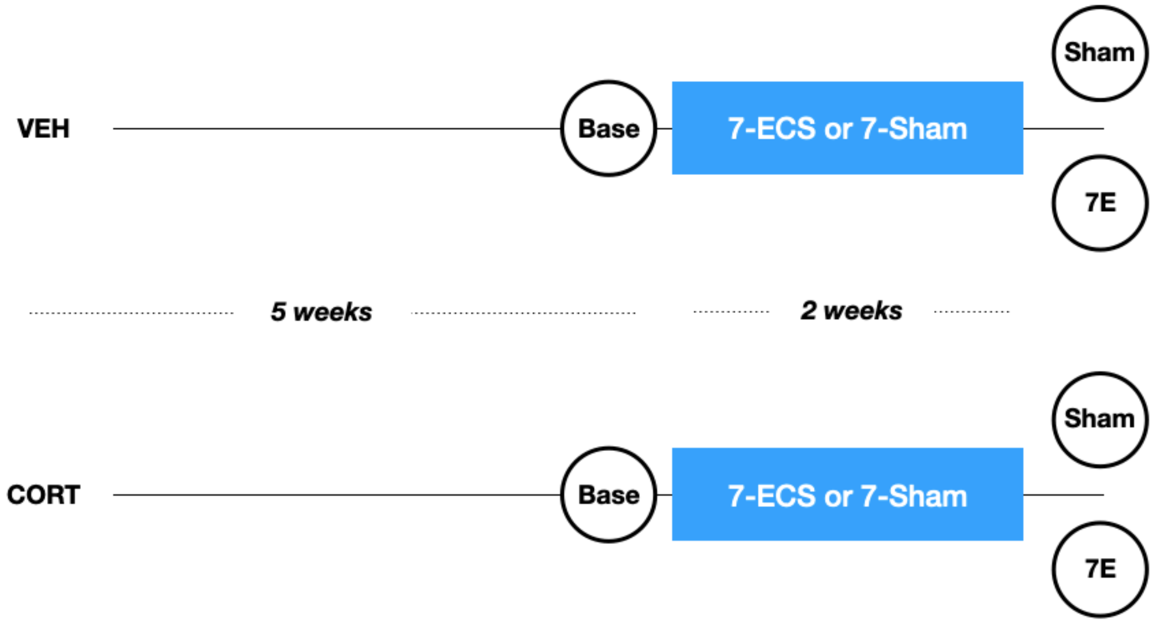


n=4 animals per group
